# Acoustic inspired brain-to-sentence decoder for logosyllabic language

**DOI:** 10.1101/2023.11.05.562313

**Authors:** Chen Feng, Lu Cao, Di Wu, En Zhang, Ting Wang, Xiaowei Jiang, Chenhao Zhou, Jinbo Chen, Hui Wu, Siyu Lin, Qiming Hou, Chin-Teng Lin, Junming Zhu, Jie Yang, Mohamad Sawan, Yue Zhang

## Abstract

Many severe neurological diseases, such as stroke and amyotrophic lateral sclerosis, can impair or destroy the ability of verbal communication. Recent advances in brain-computer interfaces (BCIs) have shown promise in restoring communication by decoding neural signals related to speech or motor activities into text. Existing research on speech neuroprosthesis has predominantly focused on alphabetic languages, leaving a significant gap of logosyllabic languages such as Mandarin Chinese which are spoken by more than 15% of the world population. Logosyllabic languages pose unique challenges to brain-to-text decoding due to extended character sets (e.g., 50,000+ for Mandarin Chinese) and complex mapping between characters and pronunciation. To address these challenges, we established a speech BCI designed for Mandarin, decoding speech-related stereoelectroencephalography (sEEG) signals into coherent sentences. We leverage the unique acoustic features of Mandarin Chinese syllables, constructing prediction models for syllable components (initials, tones, and finals), and employ a language model to resolve pronunciation to character ambiguities according to the semantic context. This method leads to a high-performance decoder with a median character accuracy of 71.00% over the full character set, demonstrating huge potentials for clinical application. To our knowledge, we are the first to report brain-to-sentence decoding for logosyllabic languages over full character set with a large intracranial electroencephalography dataset.

## Main

Numerous grave neurological conditions, including stroke and amyotrophic lateral sclerosis, may result in individuals entirely losing their capacity to verbal communicate^1,2^. Previous speech neuroprosthesis effort has demonstrated the potential to restore communication for speech-impaired patients^3^, particularly within the scope of alphabetic linguistic systems (e.g., English and Dutch), by establishing a mapping between text output and different activities related brain signals, such as hand-writing^4^, spelling^5–7^, typing^8,9^, and speech^10–14^. While such attempts have shown promise, no existing research has reported investigation on logosyllabic languages, such as Chinese and Thai language, for which the speaking population is more than 1.3 billion^15^. In contrast to the alphabetic languages, where words are constructed from combinations of limited size alphabets (e.g., 26 for English), logosyllabic languages employ logographic characters to represent either entire words or single morphemes, which cannot be typed or spelled out directly. Moreover, logosyllabic languages such as Mandarin Chinese have an extensive inventory of over 50,000 logographic characters^16,17^, each with its unique, complex glyph. For example, the English word “cat” is represented by a single character “猫” in Mandarin, and “hot” is denoted as “热”. This complexity renders the decoding of handwriting via BCIs exceedingly challenging for such languages. Considering the fact that Mandarin pronunciation is mostly detached from its glyphs, we seek a viable solution to bypass the curse of logographic systems via decoding the pronunciation of characters.

We consider *Pinyin*^18^, a widely adopted phonetic representation system based on the Latin alphabet, as the foundation for a neuroprosthesis for Mandarin speech decoding. *Pinyin* has been developed to facilitate literacy and serve as a bridge between Mandarin pronunciation and characters^19^. It adopts a single combination of initials (initial consonant of a Chinese syllable), finals (simple or compound vowel of a Chinese syllable), and tones (pitch variation at the syllable level) to multiple Mandarin Chinese characters^20^. For instance, the combination of initial ‘t’ ([tʰ]), final ‘a’ ([a]) and a high-level tone (Tone 1) can be mapped to both ‘他’ (tā, [tʰa], he) and ‘塌’ (tā, [tʰa], breakdown), which are homophones. The total number of *Pinyin* syllables (excluding variations in tone), amounts to 407. Each syllable can be combined with 4 different tones, introduces additional intricacy to speech decoding since the alterations in a tone of one syllable can lead to a significant shift in its semantic meaning^21^. Due to ambiguity in mapping *Pinyin* into characters, the surrounding context plays a pivotal role in identifying the correct character in a sentence.

We propose a Mandarin Chinese BCI system that achieves the conversion of speech-related brain signals into sentences by decoding the pronunciation of *Pinyin* syllables. Unlike some existing work, which covers only part of the vocabulary, our system can handle the full set of Chinese characters, despite being trained with only 407 syllables. This system comprises three acoustic inspired syllable element prediction models (for the initial, tone, and final, respectively) and a language model. The former identifies the pronunciation syllables corresponding to brain signals, while the latter, incorporating context and semantics, converts syllable sequence into a complete sentence with practical meanings.

Most previous studies on language prosthesis use Electrocorticography (ECoG) electrodes^6,11,13,22–24^ or Utah array electrodes^4,10,25^ to acquire cortical signals. However, some studies show that the subcortical signals have potential to enhance the effectiveness of decoder^26,27^. The current study utilized stereo-EEG to capture both cortical and subcortical neural signals simultaneously from four participants (two males and two females, all young native Mandarin speakers) undergoing intracranial monitoring for epilepsy (Fig. 1a). For training the model, participants were instructed to read 407 monosyllabic characters, which correspond to the 407 distinct *Pinyin* syllables (the upper panel in Fig. 1b). These 407 syllables are specially designed to contain all the initial and final pairs, so the pronunciation of almost all Mandarin Chinese characters is included. Additionally, the participants read 100 Mandarin sentences varying in length from two to more than ten characters (the lower panel of Fig. 1b), for evaluating the performance current speech BCI.

**Fig. 1.**
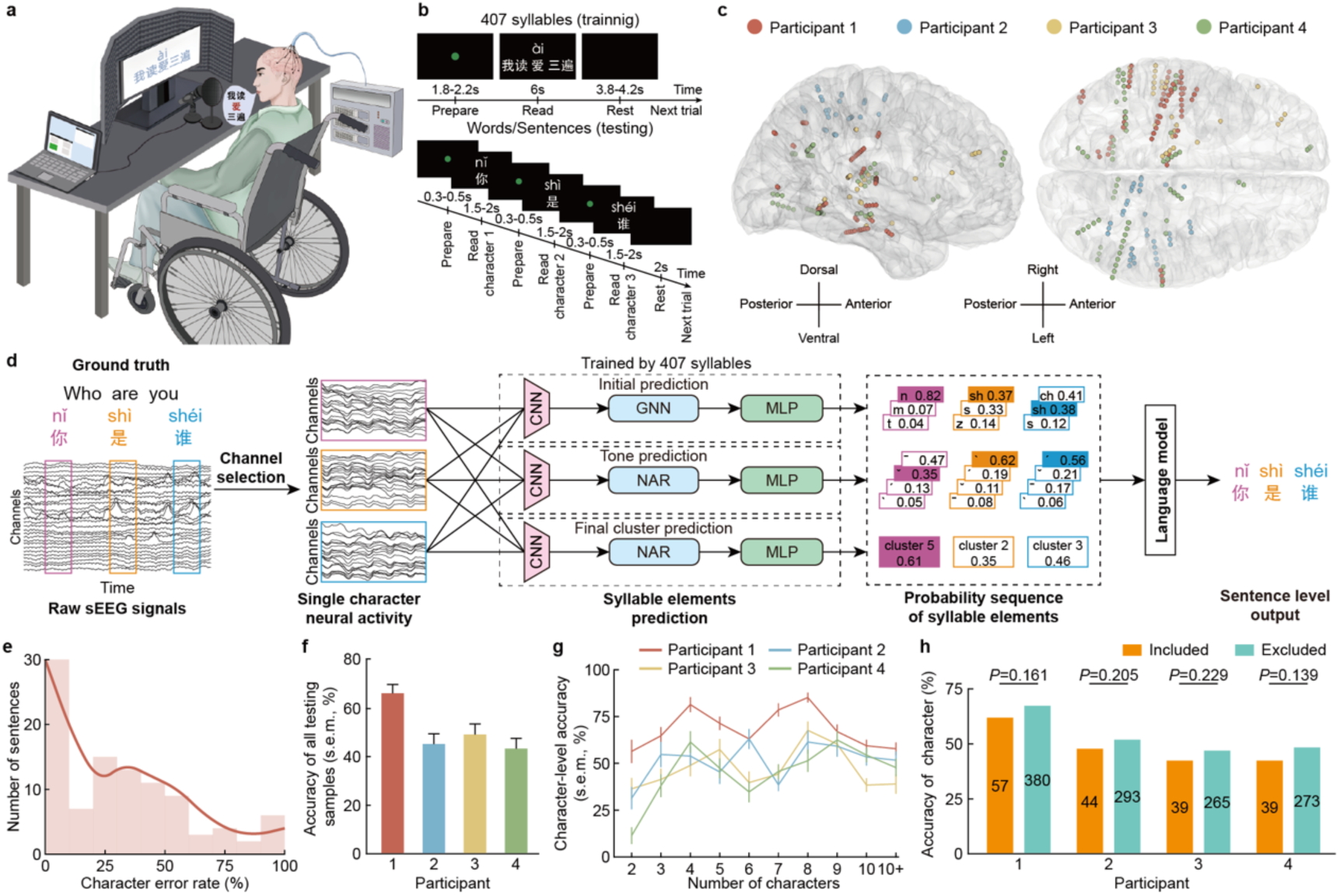
| The pipeline and performance of the brain-to-sentence decoder designed for logosyllabic language decoding. **a**, After the electrodes were implanted, participants took part in the study on the wheelchair, speaking aloud the content displayed on a screen. Audio data was collected using a directional microphone, and neural signals were simultaneously recorded using a multi-channel electrophysiological recording system. **b,** Two tasks designed for collecting training used data (upper panel, 407 *Pinyin* syllables cover almost all Mandarin Chinese characters) and evaluating used data (lower panel, 100 sentences ranging from 2 to over 10 characters in length). **c,** The distribution of selected decoding channels across participants is illustrated on the standard Montreal Neurological Institute (MNI) template brain. Each participant’s channels are denoted by distinct colors, with directional indicators provided for clarity. **d,** A three-phase decoder, including channel selection, syllable elements prediction and language model correction, is designed to output whole Mandarin Chinese sentences by decoding speech related sEEG signals. **e,** The distribution of decoding accuracy of 100 test samples (Data from participant 1). Outlines are kernel density estimates of the distributions. **f,** The decoding accuracy (mean ± s.e.m.) of all evaluating samples, which is the average of the proportion of correct characters in each sample. **g,** The character-level accuracy (mean ± s.e.m.) of sentences composed of different lengths. **h,** The comparison of decoding accuracy of characters appeared in the training reading corpus (Included) and those absent (Excluded) with the number of correctly decoded characters displayed on the corresponding bar. The significance was calculated by bootstrapping.

Trained using such data, the BCI system achieved the direct decoding of brain activity directly into text sentence of logosyllabic language, where the sentences are highly unconstrained and free from the limitations imposed by the training data, potentially enabling the reconstruction of free language expression in patients who have lost the ability of verbal communication. Furthermore, we found that the performance of this acoustic-inspired decoder is driven by both cortical and subcortical signals.

## Result

### Overview of decoder design

For each participant, we used signal from the selected channels associated with speech and excluded those located in visual cortex and white matter (Fig. 1c). A three-phase (channel selection, syllable elements prediction and language model) decoder were proposed for decoding the neural signal to the complete sentences (Fig. 1d). During the syllable elements prediction phase, three convolutional neural networks (CNNs) were trained to extract features representing the initials, tones and finals from the preprocessed sEEG signal segments of the syllables being spoken. For each syllable, we aligned the tone and final features with acoustic features extracted from the corresponding audio signal using neural-audio regularization (NAR). Initial features were mapped into articulatory features, with their relationships further refined by a graph neural network (GNN). In the evaluating process of decoding complete sentences, we used the above three trained models to predict initial, tone, and final syllable by syllable, and finally formed a probability sequence. In the third phase, a language model was established to output the most probable sentence based on the probability sequence of predicted elements of all syllables. The language model was trained on encyclopedia-style question-answer dialogues retrieved from Common Crawl. The probability sequence of top 3 initials, top 3 tones and the leading final cluster (one final cluster contains several finals with similar acoustic characteristics) form the output of the three syllable element prediction models. Furthermore, the language model is capable of translating *Pinyin* sequences into Mandarin sentences, correcting errors in syllable element predictions, and enhancing the overall performance of sentence outputs.

### Performance of the brain-to-sentence decoder

To fully illustrate the decoding capability of this system in the natural Mandarin environment, we randomly selected sentences for daily use ranging in length from 2 characters to more than 10 characters, taking 10 of each length for a total of 100 test samples. First, we analyzed the distribution of the average character error rate for each sentence (Fig. 1e). Supplementary Tab. 1 presents examples of decoded sentences across various character error rate levels. For participant 1, the median character accuracy is 71.00%, with 30.00% of the decoded sentences being completely accurate (Fig. 1f). The decoded sentences exhibit a low error rate and high intelligibility, making them both meaningful and viable for practical applications.

We calculated the character-level decoding accuracy and found that the average accuracies for the four participants were 66.62%, 51.37%, 46.34%, and 47.56%, respectively (Fig. 1g). Notably, sentences of moderate length (about 4-8 characters) yielded higher accuracies. Conversely, the accuracy for decoding shorter phrases of 2-3 characters was lower, primarily due to the lack of sufficient linguistic context. This limitation hindered the model’s ability to distinguish between phrases with identical pronunciations but different meanings, such as ‘前沿’ and ‘前言’, with the same *Pinyin* syllable (qián yán) and different meanings (frontier and preface, respectively). We verify that the generalization performance of the model is strong, and it can solve the parts not covered by the training data in tens of thousands of Mandarin Chinese characters (Fig. 1h), which also highlights the effectiveness of our acoustic inspired decoding strategy.

### Design and performance of initial prediction model

In the Mandarin Chinese *Pinyin* system, the initial consonant is termed initials, and there are a total of 21 distinct initials. The articulation of initials is produced by overcoming the obstruction of airflow as it passes through the oral or nasal cavity. Four features, place of articulation (POA, including bilabial, labiodental, alveolar, retroflex, alveolo-palatal and velar), manner of articulation (MOA, including plosive, affricate, fricative, nasal and lateral), devoice (including voiced and voiceless) and aspiration (including unaspirated and aspirated) uniquely describe the articulation process of each Mandarin initial consonants (Supplementary Table 2). Figure 2a shows the articulation process and corresponding articulation features of two initials, which is ‘t’ ([tʰ]) and ‘m’ ([m]).

**Fig 2.**
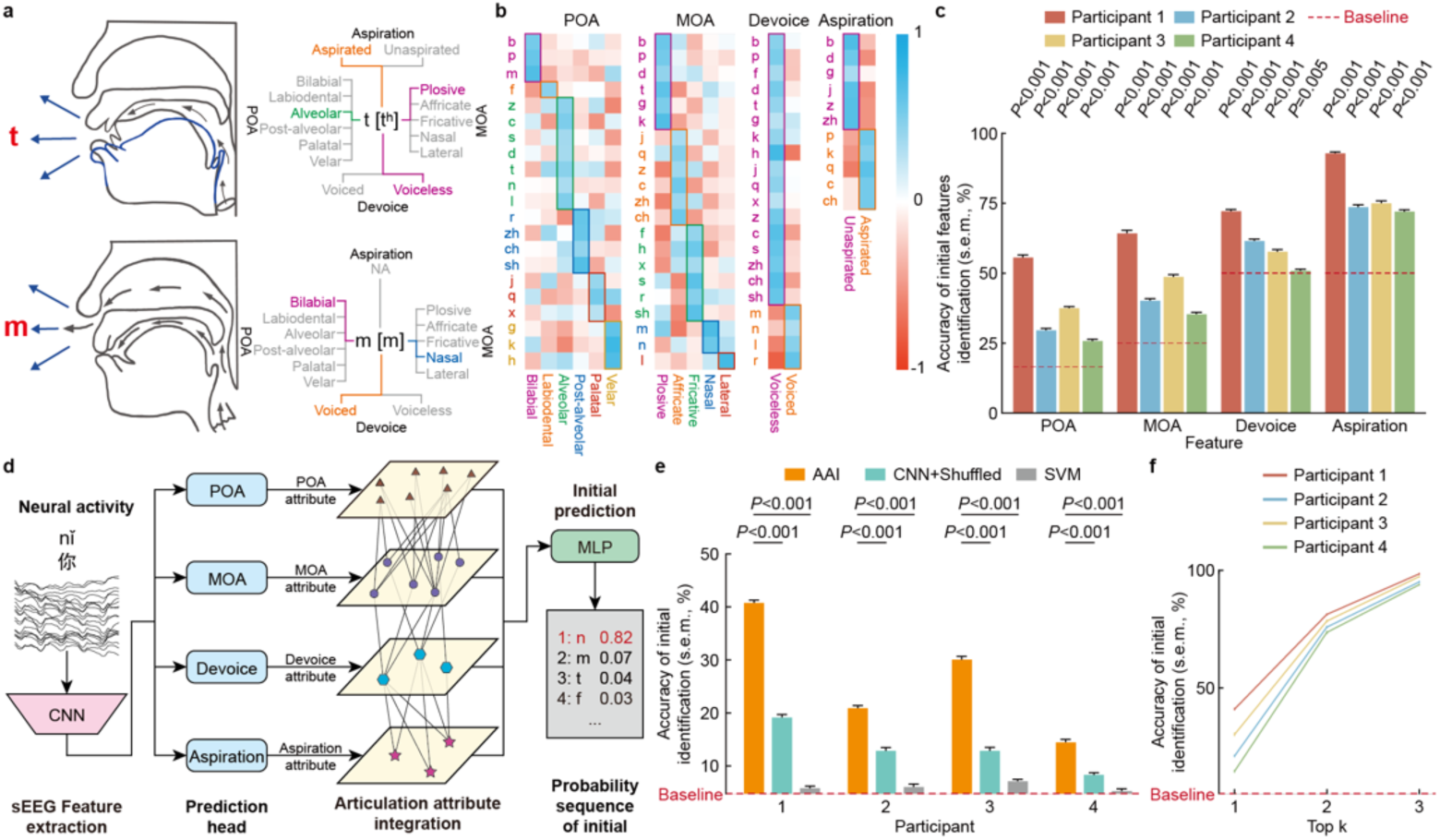
| Design motivation and performance of initial prediction model. **a**, Articulatory descriptions of the initials ’t’ and ’m’ characterize acoustic-phonetic features (on the left). Within the dynamic articulation process, the gray section denotes the occlusive phase of articulation, while the blue section indicates the release phase. The multiple acoustic-phonetic features are combined to describe these two phonemes (on the right). **b,** The correlation between specific initials and the average of their corresponding category is visualized through a series of heat maps, where the color representing each initial is matched to the color of its category. **c,** Prediction accuracy (mean ± s.e.m.) of four initial features for four participants. **d,** The initial prediction model utilizes a Graph Neural Network (GNN) to integrate various acoustic-phonetic features. This Articulation Attribute Integration (AAI) approach effectively captures the complex interactions of voicing features to accurately map neural signals onto these feature spaces. **e,** The prediction accuracy (mean ± s.e.m.) of initial was measured for four participants (AAI) and compared with scenarios where acoustic-phonetic feature labels were shuffled (CNN + Shuffled) and against the performance obtained from the support vector machine (SVM). **f,** The probabilities (mean ± s.e.m.) of the correct initial being included in the top 1, top 2, and top 3 probability sequence.

To assess the similarity in neural activity during the articulation of initials sharing common features, we analyzed the correlation between specific initials and the average of their corresponding category (Fig. 2b). Our findings revealed a positive correlation for place of articulation (POA: r = 0.52), manner of articulation (MOA: r = 0.54), devoicing (r = 0.49), and aspiration (r = 0.59). This correlation is visually represented by four diagonal lines across the heatmap for each feature. Moreover, the correlation is significantly higher than that with irrelevant articulation features (*P* < 0.001 for POA, MOA, devoice and aspiration). indicating that the sEEG signals captured distinct information related to categorical articulation features. To further explore this, we developed classification models to decode these four articulatory features from the sEEG signals of each initial’s trial (Fig. 2c). The classifiers demonstrated that the sEEG signals contained significant information about these features, with decoding accuracies substantially surpassing the chance level for all participants.

Building upon these insights, we build the initial prediction model by identifying the acoustic-phonetic features (Fig. 2d, details in method). We first map the neural signals to representations in the acoustic-phonetic feature spaces of POA, MOA, aspiration and devoice. Given that the voicing of initials results from the interactions among diverse voicing features, we address this inherent relationship with a GNN as an articulation attribute integration (AAI). As a result, the accuracy rates from the model are significantly above the chance level for the four participants (*P* < 0.001, Fig. 2e), achieving rates of 41.07%, 21.10%, 30.32%, and 14.65%, respectively, where the chance level is 5%. However, the sole prediction result from the model is evidently insufficient in accuracy for input into the language model for sentence transformation. Therefore, we calculated the accuracy considering multiple candidate outputs from the model. Notably, there was a significant increase in prediction accuracy as the number of candidate results grew (*P* < 0.001). In the top 3 most probable predictions, the probability of containing the correct initial for the four participants was 99.24%, 97.12%, 98.39%, and 95.03%, respectively (Fig. 2f). In summary, the initial prediction model can identify accurate initial results in most trials based on sEEG signals from selected channels, and its near-perfect top-3 accuracy narrows the possible initial candidates from 21 down to 3. This reduces the computational burden on the third phase of the decoder and enhances the efficiency of the selection process of language model.

### Design and performance of tone prediction model

Tone, a crucial aspect of Mandarin syllables, differentiates both lexical and grammatical meanings through variations in pitch carried by the finals^21,28^. We visualized the pitch changes over time for the four Mandarin tones (Fig. 3a), highlighting their distinct patterns and stability across trials of the same tone. To distinguish these tones quantitatively based on pitch variations, we employed a support vector machine (SVM) classifier trained on pitch data from all trials, regardless of the rhymes on which the tones were borne. The four different tones can be distinguished from the perspective of pitch variations, and the values on the diagonal are significantly higher than the off-diagonal values in Fig. 3b (*P* = 0.017). The final classification accuracy was 57.51%, significantly above the chance level (*P* < 0.001). Consequently, we incorporated pitch into the training process of the tone prediction model (Fig. 3h, details in method). Aligning the distribution of sEEG data in the neural space with the distribution of audio data, particularly concerning pitch features, enhances the extraction of neural features for tone categorization. We incorporate a neural-audio regularization (NAR) term based on similarity measurement in addition to the categorical cross-entropy loss during model training. Audio signals will solely be utilized in the decoder training process and not during evaluating.

**Fig. 3.**
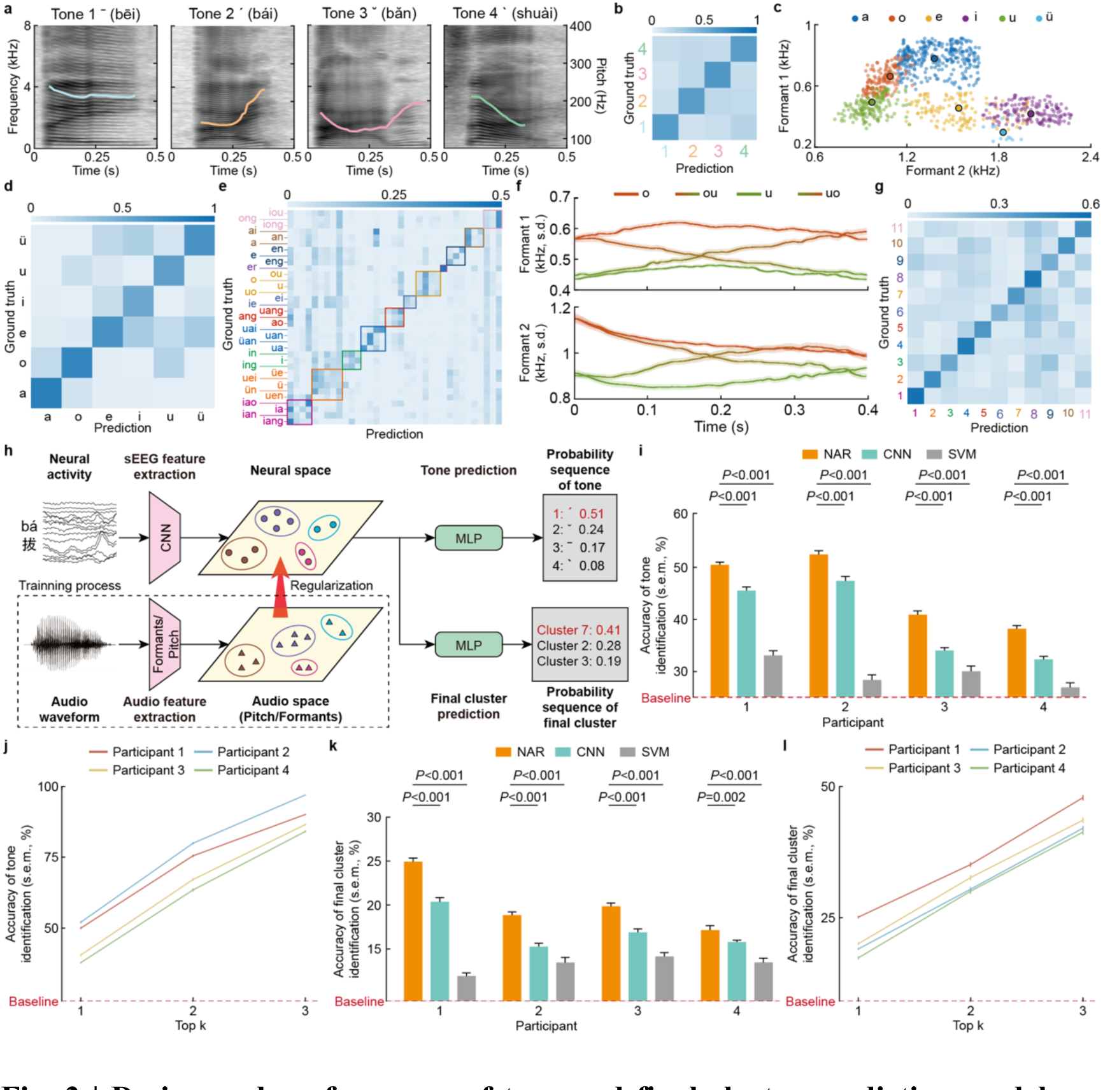
| Design and performance of tone and final cluster prediction models. **a**, Examples of the display of Mel-frequency spectrograms and pitch curves for the four tones, with data sourced from participant 1. **b,** Based on pitch feature extracted from audio signal, a SVM model was used to classify the four tones (participant 1). **c,** The visualization results of the Formant features for six single finals are shown. Since the formants are relatively stable during the articulation of single finals, the average value of the entire process is used to represent the corresponding formant feature. Dots of the same color represent different samples of the same single final, with the larger dot in the corresponding color representing the average of all the dots. **d,** Based on formant features extracted from audio data, a SVM model was used to classify the six single finals (participant 1). **e,** Based on formant features extracted from audio data, a SVM model was used to classify the 35 finals (participant 1). Finals of the same color were clustered into the same category using the k-means clustering method. **f,** The dynamic changes of Formant 1 and Formant 2 during the articulation process of four phonetically similar finals (‘o’, ‘u’, ‘ou’, and ‘uo’) are depicted (mean ± s.e.m.). **g,** Based on formant features extracted from audio data, a SVM model was used to classify the eleven final clusters (participant 1). **h,** The model design for tone and final cluster prediction. This approach aligns the distribution of sEEG data in the neural space with the audio data distribution, especially in terms of pitch and formant features, to improve the extraction of neural features for tone and final cluster categorization. Particularly, we employed a neural-audio regularization (NAR) term, based on similarity measurements, alongside the categorical cross-entropy loss during the training of the model. **i,** The prediction accuracy of tone was measured for four participants and compared with scenarios where acoustic-phonetic feature labels were shuffled (Shuffled) and against a baseline of accuracy (Chance). **j,** The probabilities (means ± s.e.m.) of the correct tone being included in the top 1, top 2, and top 3 prediction results for each of the four participants. **k,** The prediction accuracy of final cluster was measured for four participants and compared with scenarios where acoustic-phonetic feature labels were shuffled (Shuffled) and against a baseline of accuracy (Chance). **l,** The probabilities (means ± s.e.m.) of the correct final cluster being included in the top 1, top 2, and top 3 prediction results.

The incorporation of NAR significantly enhanced the model’s performance, yielding prediction accuracies of 50.60%, 52.75%, 41.19%, and 38.48% for the four participants, respectively, all markedly above the chance level (*P* < 0.001 for all, as shown in Fig. 3i). These accuracies also surpassed those of models trained without NAR, highlighting NAR’s importance. Given that a single prediction may not capture all correct tones, we adopted a strategy of selecting the top 3 most probable tones for each prediction. This adjustment significantly improved the accuracy of the final sentence predictions, with the correct tones being included in the top 3 predictions with probabilities of 90.77%, 97.66%, 87.21%, and 84.76% for the participants, respectively (Fig. 3j). Importantly, this probabilistic information about the tones is also conveyed to the language model, providing it with richer reference data for more accurate sentence generation.

### Design and performance of final prediction model

Mandarin syllables comprise 6 simple and 29 compound finals, whose articulation involves vocal cord vibration and nuanced tongue positioning. The frequency spectrum’s formant features, F1 and F2, correspond to the tongue’s vertical and anteroposterior positions, respectively. We visualized the formant features F1 and F2 for all trials of six simple finals, revealing a clear clustering effect (Fig. 3c). An SVM classification model trained on F1 and F2 successfully differentiated the audio of the 6 simple finals, achieving a classification accuracy of 66.35%, significantly surpassing the chance level (*P* < 0.001Fig. 3d).

However, once the compound finals were incorporated, the classification results deteriorated (15.46%, compared to the chance level, *P* = 0.14, Fig. 3e). This decline in performance can be attributed to the dynamic nature of compound finals articulation, which involves transitioning from one articulatory shape to another. For instance, we visualized the changes in F1 and F2 during the articulation of the simple finals ’o’ and ’u’ alongside the compound finals ’ou’ and ’uo’ (Fig. 3f). The compound final ’ou’ is articulated by transitioning from the sound of ’o’ to ’u’, while ’uo’ undergoes the opposite transition, making their articulatory configurations exceedingly similar and hence challenging to distinguish. Therefore, based on the resonance peaks of the finals in the audio, we employed the k-means algorithm to cluster finals with similar articulatory configurations, ultimately forming 11 distinct finals clusters (Fig. 3e). The SVM classification models were used to identify the final clusters, achieving an accuracy of 40.86%, which is significantly higher than the chance level (*P* = 0.013, Fig. 3g).

Extracting the formant features from audio has proven beneficial for classifying final clusters. Therefore, we also employed the model used to predict tones with NAR to predict the final clusters based on sEEG signals (Fig. 3h, details in Method). Despite the finals’ complexity, the model demonstrated accuracies of 25.10%, 19.02%, 20.04%, and 17.33% across participants, all statistically significant against the chance level. As the number of candidate clusters increased, the probability of including the correct final also rose (Fig. 3i). The inclusion of multiple finals within a single predicted cluster aimed to reduce the selection burden in the decoder’s third phase, acknowledging that the precision of initials and tones predominantly conveys the necessary information for accurate sentence prediction in Mandarin, thus mitigating the impact of less accurate final predictions.

### Contributions of different brain regions

In this study, we attempted to independently utilize signals from cortical and subcortical brain regions to predict three syllable elements. Comparative results at the element level were consistent, yet notable variances at the individual participant level emerged (Fig. 4a), indicating that the decoding efficacy is highly dependent on the specific brain regions engaged and the electrode placement. Participant 1 showed enhanced decoding performance with cortical signals, whereas Participant 3 demonstrated better outcomes with subcortical signals. The results for Participant 4 indicated no significant difference between the decoding effectiveness of the signals from these two regions. Overall, this suggests that while cortical regions are influential in speech decoding, the contribution from subcortical regions is also meaningful and should not be overlooked.

**Fig. 4.**
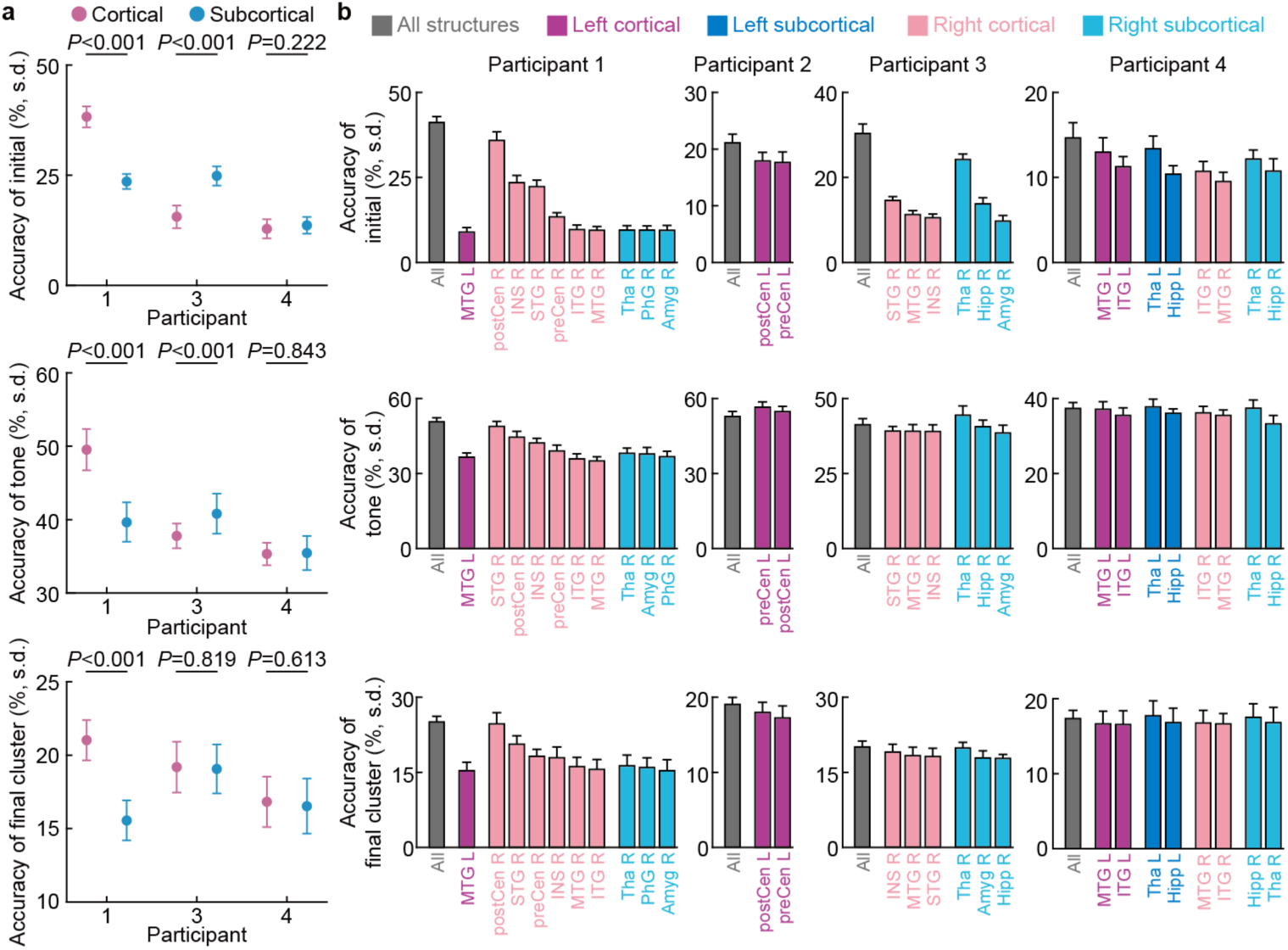
| The decoding accuracy and contributions of each anatomical area. **a**, The prediction accuracy of initial, tone and final cluster using neural signals from cortical and subcortical brain structures. Accuracy data are means ± S.D., and the results were presented for only 3 participants (1, 3 and 4), as no chosen electrodes of participant 2 were in the subcortical brain structures. **b,** The prediction accuracy (means ± S.D.) of initial, tone and final cluster using neural signals from different anatomical areas in cortical and subcortical structures of right and left hemispheres of the brain. The regions of the brain where the electrodes were located were different for four participants. Only the results of some important language-related anatomical area are selected to show. STG: superior temporal gyrus, MTG: middle temporal gyrus, ITG: inferior temporal gyrus, postCen: postcentral gyrus, preCen: precentral gyrus, INS: insula, Tha: thalamus, Hipp: hippocampus, Amyg: amygdala, PhG: parahippocampal gyrus.

To further visualize speech-related information encoded in neural signals from different brain regions, we used signals from the electrodes located in the same regions as input to separately predict the initial, tone, and final cluster. Figure 4b shows the prediction accuracy of three syllable elements from neural signals in different brain regions for four participants. Due to the variation in the brain regions and the number of electrodes implanted in different participants, cross-participant comparisons are not feasible. Results from Patient 1 indicate that within the cortical areas, the Superior Temporal Gyrus (STG), Insula (INS), and the ventral Sensorimotor Cortex (composed of the postcentral gyrus and precentral gyrus, abbreviated as vSMC), exhibit higher decoding accuracies. This observation aligns with traditional theories of brain region function in speech production^20,21^, and speech perception^22,23,29–31^. Within the subcortical areas, we observed that the thalamus exhibited significantly higher accuracy in certain prediction tasks (such as the initial prediction for participant 3) compared to other brain regions. Previous research has reported the involvement of the thalamus in human language processing^32,33^. Our findings further reveal that the thalamus is also engaged in human speech processing.

### Language models improve decoding accuracy

In this study, the language model is tasked with selecting the appropriate components to form pinyin syllables from the probability sequences of the three syllable elements (initials, tones, and finals). Subsequently, it was required to identify the most appropriate Mandarin Chinese characters matching each pinyin syllable’s pronunciation, thereby constructing sentences that are both semantically and grammatically coherent.

To enhance the accuracy of this process, we determined that it was necessary to input the top 3 initials, top 3 tones, and the first final cluster as element possibility sequences into the language model in the decoder. We calculated the accuracy of the syllable elements converted from the final output sentences (element-level accuracy), where each element is selected by the language model from the probability sequences. The element-level accuracy for the four participants (Fig. 5a) are 80.98%, 66.82%, 64.98%, and 63.80%, respectively, which are significantly higher than the top 1 accuracy of the three element prediction models (Fig. 5b). From the aforementioned results, it is evident that the language model plays a vital role in selecting the correct elements and correcting erroneous ones, thereby mitigating the accuracy demands placed on the brain signal decoding component.

**Fig. 5.**
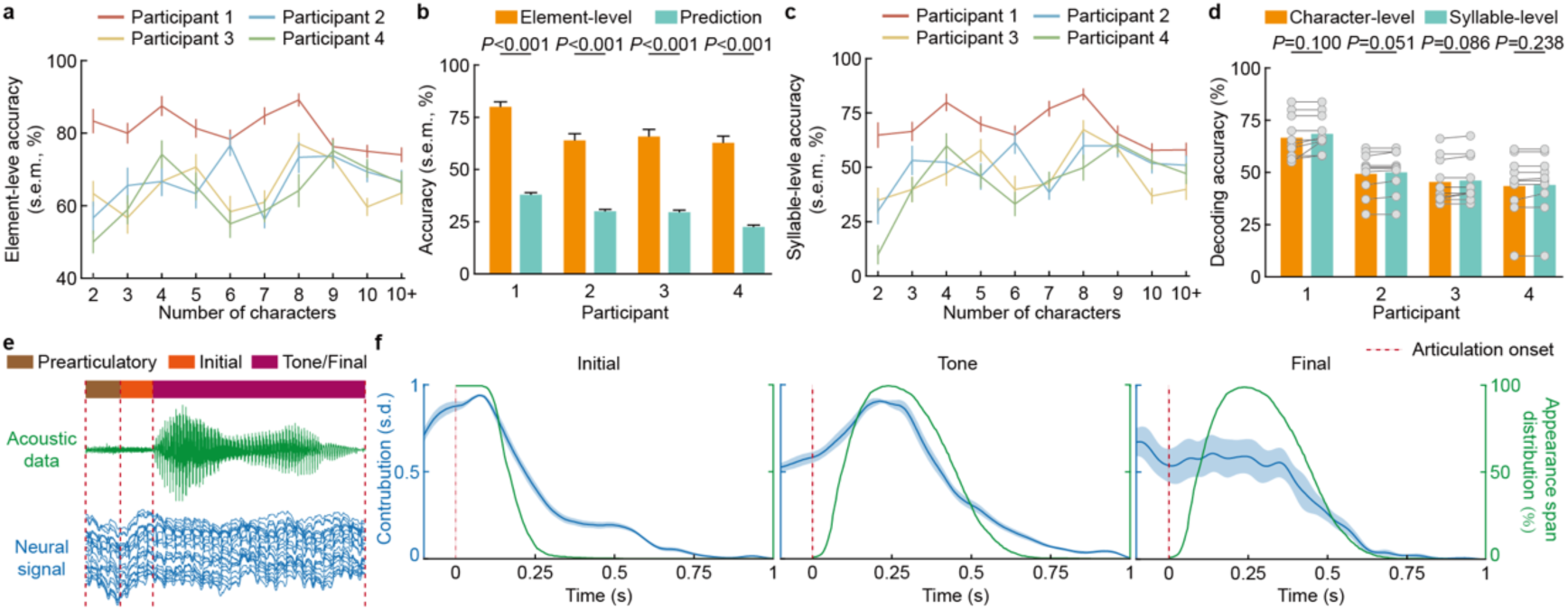
| The analysis of language model and three syllable element prediction models. **a**, The element-level accuracy (mean ± s.e.m.) of sentences composed of different lengths for four participants. **b,** The comparison between element-level accuracy (mean ± s.e.m.) and the accuracy (mean ± s.e.m.) of three element prediction models decoding along (Prediction). **c,** The syllable-level accuracy (mean ± s.e.m.) of sentences composed of different lengths for four participants. **d,** The comparison between character-level accuracy (mean ± s.e.m.) and syllable-level accuracy (mean ± s.e.m.). The gray dots represent the accuracies of sentences in different lengths (ranging from 2 to 10+ characters, with a total of 10 different conditions), where the significance comparison is conducted using paired t-tests. **e,** Acoustic data and neural signal of articulation process of fěn. The articulation process is divided into three periods by phonetics experts. The first is the prearticulatory period (100ms). The second is the initials articulation period, which is generally shorter in length. The last is the articulation process of tones and finals. Because the tones are carried on the finals, the pronunciation process overlaps. According to the theory of phonetics, the articulation process of initials and finals overlap. This study divides them according to the average position in the overlapping part. **f,** The temporal saliency on neural signals of three syllable element prediction models used to identify the initial, tone, and final clusters, respectively, over the entire duration of a syllable (Blue line, mean ± s.d.). The green line indicates the appearance span distribution with respect to time for the initial, tone, and final of all 407 syllables, with the start and end points marked by experienced phoneticians.

Furthermore, to evaluate the language model’s proficiency in transforming syllable sequences into accurate character sequences through semantic integration, we computed the syllable-level accuracy and compared it with the character-level accuracy. The syllable-level accuracies for the four participants were 68.93%, 50.52%, 46.68%, and 44.50%, respectively (Fig. 5c), which closely aligned with the character-level accuracies (Fig. 5d). This convergence indicates that the language model adeptly translated nearly all correct syllables into their accurate character counterparts. The few observed instances of homophone conversion errors, such as ‘他’ (tā, [tʰa], he) and ‘她’ (tā, [tʰa], she), did not detract from the overall comprehension of the sentences for native Mandarin speakers. The ability of the language model to convert pinyin and integrate context plays a decisive role in accurately translating syllable sequences, enabling it to select the correct characters from a vast array of homophones.

### Model attention across time

Acknowledging the challenges posed by the brief and often ambiguous transitions between the initial and final parts of syllables (Fig. 5e), our study approaches syllable decoding by treating the neural signals of the entire articulation process, including a 100ms prearticulatory period, as an integrated unit. This method allows us to preserve crucial information that might otherwise be lost during segmentation. We investigate if the temporal focus on the brain signals of the syllable components (initial, tone, and final) decoding models align with the vocalization processes of these components as reflected in the corresponding speech signals by calculating the temporal saliency of the syllable element prediction models.

Our analysis, illustrated in Figure 5f (first panel for initial and mid panel for tone), shows a strong correlation between the temporal saliency on brain signals and appearance span distribution with respect to time of these syllable components in terms of audio data. This suggests that our models can effectively identify and capture syllable element relevant brain signal segments throughout the articulation process for accurate prediction. Compared to the initial and tone prediction models with prominent temporal focus regions, we observed a more uniform distribution in the saliency distribution for the finals prediction model (Fig. 5f, last panel). This indicates that the finals prediction model struggles to pinpoint final-related brain signal segments, which in turn, might explain the model’s comparatively lower performance in predicting finals accurately.

### Channel contribution

In the first phase of the decoder, the signals from all regions related to speech are utilized for language decoding. To determine their contribution in the trained model, we compute a saliency score for each channel as indication of contribution for syllable components prediction (Supplementary Figure 1). Subsequently, we also perform a correlation analysis with the decoding accuracy obtained using channels from individual brain regions as input data. The correlation between the prediction accuracy for three syllable elements and their contribution in three prediction models were found to be significant across all four patients (Supplementary Figure 2).

## Discussion

We tackled the challenges posed by the complexity of logosyllabic languages in speech BCI by taking the speech of individual character syllables as the decoding unit, achieving the transformation of speech-related brain signals into sentences for Mandarin Chinese with a promising performance. We show that a training dataset that includes all 407 valid syllables in Mandarin, suffices for decoding the full Chinese character set, when combined with a suitable language model. Additionally, we used monosyllable with tone as the modeling unit because previous unit selection study has proved that the use of syllable with tone effectively increases the speech recognition accuracy of Mandarin Chinese compared to other units such as characters or phonemes^34^. Since around 60∼70% of the world’s languages are tonal^35^, which is not reflected in the mainstream research on speech BCIs, our study of Mandarin Chinese contributed fundamental insights to the broader landscape of tonal language interface design and brain-computer system development.

Recently, numerous language neural prosthetic systems have incorporated language models as a component of their architecture^10,11^. In this study, we have developed a language model suitable for Mandarin Chinese speech BCI systems by integrating *Pinyin* input with an N-gram model^36^. The capability of the language model for identifying grammatical Mandarin Chinese text, resolves the ambiguity in pronunciation to character mapping, and for the *Pinyin* predictor, which in turn significantly improves the accuracy of the final sentence prediction. While our language model is intended for open and unrestricted Mandarin Chinese language communication, it is worth noting that in specific environments and scenarios, such as in the domestic settings of individuals with disabilities, the language model can be trained on a restricted, predefined corpus to yield more accurate output results.

Our decision to exclude the decoding of neutral tones was based on the observation that, aside from a few particles, the majority of neutral tones evolve contextually from their corresponding full tone counterparts^37^. This evolution alters their phonetic characteristics depending on the preceding tones in spoken language but remains unchanged in the written form employed in our research^38^. Nonetheless, future investigations might delve further into the nuances of neutral tones for a more comprehensive understanding of their role in Mandarin Chinese BCI research.

We observed notable variations in the decoding performance when distinguishing between single finals, compound finals, and nasal finals, with the latter two showing less favorable results. This may be attributed to the finer timing and position properties of articulatory gestures during a continuous shift of vowels and nasals in compound finals and nasal finals compared to single finals^39^. Thus, we implemented a k-means clustering approach, which uses F1 and F2 frequency parameters to classify these non-single finals into 11 clusters. These clusters share similar acoustic properties and articulatory patterns, such as ’in’ and ’ing,’ grouped based on the nucleus in the structure of finals, as the nucleus bears highest sonority scale^40^, for simplifying the decoding process. These can provide a foundation for further investigations into the complexities of decoding compound finals and nasal finals in the context of Mandarin Chinese BCI systems.

Addressing the challenges of mitigating the need for extensive data acquisition to facilitate cross-day model transfer was beyond the immediate scope of this study. Nonetheless, this endeavor stands as a pivotal requirement in the development of a genuinely pragmatic and impactful system. Furthermore, it is essential to verify the online decoding performance, despite the promising outcomes of our offline analyses. Recent large language models benefit from extensive training data^41^, but combining data across subjects and time points for neurophysiological signals does not yield commensurate enhancements due to inherent cross-participant and cross-day variations. Considering the model adaptation in a lightweight manner to ensure cross-day accuracy and enhancing decoder performance through the use of multi-subject data, the critical endeavor of optimizing the efficient utilization of data from diverse days and individuals remains a key focus for forthcoming research.

## METHODS

### Participants and task

#### Study participants

This study enrolled four participants—two females and two males, aged 20, 25, 19, and 19 years for Participants 1, 2, 3, and 4, respectively. They were undergoing epilepsy treatment at the Second Affiliated Hospital of Zhejiang University School of Medicine. As part of their treatment, they had sEEG electrodes surgically implanted (with the number of electrodes being 12, 8, 11, and 13 for Participants 1, 2, 3, and 4, respectively) to monitor epileptic seizures for 1-2 weeks. All participants were native Mandarin Chinese speakers with the left hemisphere identified as the dominant hemisphere for language processing. Prior to data collection, all patients were informed of study procedure and signed informed consent form to participate. The study was approved by the Ethics Committee of the Second Affiliated Hospital of Zhejiang University School of Medicine.

#### Task design

The design of this Mandarin Chinese Brain-Computer Interface (BCI) system is predicated on decoding the neural signals during the articulation process of Mandarin Chinese characters, ultimately converting them into Mandarin Chinese sentences. Hence, the tasks were specifically tailored for this purpose. We designed two distinct sets of tasks for training and evaluating the BCI system respectively. The first set of tasks involves reading 407 unique Mandarin Chinese characters aloud. Although there are only 407 characters, they encompass all the phonetic syllables of Mandarin Chinese characters. The objective is to enable the system to decode virtually all Mandarin Chinese characters with as minimal training as possible, achieving a truly unrestricted Mandarin Chinese decoder. The second set of tasks, aimed at assessing the performance of the BCI system, employs Mandarin Chinese sentences of varying lengths as evaluating samples, to closely simulate the performance of this system in everyday life scenarios.

### Task for collecting training data

During the model training data collection phase (as depicted in Fig 1b, upper panel), participants were required to read 407 Mandarin Chinese characters corresponding to Mandarin Chinese *Pinyin* syllables with tones, three to five times (3 to 5 trials per syllable × 407 syllables, depending on the monitoring duration for different participants). These 407 syllables encompass virtually all the pronunciations of Mandarin Chinese characters. Moreover, we designed the study to maintain as balanced a number of syllables as possible across the four tones, as shown in Supplementary Figure 2. To mimic natural speech patterns closely, each character was embedded within a carrier sentence by adding fixed characters before and after the target character. Thus, in every trial, participants read a complete sentence incorporating the designated syllable. For example, for the target syllable “ài,” the constructed sentence for reading was “我读 爱 三遍” ([ài], translating to “I read love three times”). All participants were pre-acquainted with the reading materials.

### Task for collecting evaluating data

During the data collection process for evaluating the model (as illustrated in Fig 1b, lower panel), participants were required to read a total of 100 common Mandarin Chinese sentences, with each word or sentence being read only once. These 100 samples were divided into 10 groups based on character length, ranging from 2 characters to more than 10 characters, with each group containing 10 different samples of the same character length. Participants were instructed to read aloud character by character at a normal pace, following prompts displayed on a monitor in front of them. Each character was allotted a reading time of 1.5 to 2 seconds, with a preparatory period of 0.3 to 0.5 seconds before reading each character. All evaluating data were collected within a single day.

### Neural signal

#### Electrode anatomy localization and visualization

We utilized SPM12 for co-registering each participant’s preoperative T1 MRI with their postoperative CT (which includes electrode locations). Subsequently, we employed the FreeSurfer neuroimaging analysis software^42^ to reconstruct the pial surface and determine the anatomical structure where each contact (channel) is located. The standard Montreal Neurological Institute template brain^43^ was used for visualization in Fig. 1c.

#### Neural signal recoding

The electrode implantation plan, encompassing the location and number of implants, was devised solely for the treatment of epilepsy. Neural signals were recorded using a multi-channel electrophysiological recording device, specifically the Neurofax EEG-1200 produced by Nihon Kohden Corporation, Japan, and were recorded at a sampling rate of 2000Hz.

#### Preprocessing of neural signal

The neural signals collected were anti-aliased (low-pass filtered at 200 Hz) and notch filtered at 50 Hz, 100 Hz, 150 Hz, and 200 Hz to eliminate line noise, and channels exhibiting poor quality were excluded. Poor signals were defined as channels with a low signal-to-noise ratio or abnormal recordings identified during the inspection process. And downsampling was not performed during preprocessing. All neural signals were segmented according to task annotations, aligning them with the pronunciation process of individual characters.

#### Channel selection

To find speech-related channels, we compared the power spectral density (PSD, computed using Welch’s method) of neural signals from speech and silence parts within the same trial. For each channel (without channels located in white matter), a paired t-test was conducted to compare the mean PSD values of the speech condition (after speech onset) and silence condition (before speech onset). All resultant p-values were corrected to control the false discovery rate (FDR) using the Benjamini-Hochberg procedure^44^. Finally, channels exhibiting a significant difference (*P* < 0.05) between speech and non-speech conditions were selected. In addition, based on the reconstruction and localization results of all electrodes, certain channels located in the visual cortex (such as those situated in the occipital lobe) were also excluded. This exclusion was implemented to prevent visual-related signals from contributing to the decoding process. The locations of the channels ultimately selected for participation in the decoding process are displayed in Fig. 1c, with the channels of the four participants represented in different colors.

### Audio signal

#### Audio recording

Microphone recordings were obtained synchronously with the neural recordings using a directional microphone (manufactured by Audio-Technica) at 44.1kHz. Given that the recording environment was situated within a hospital ward, temporary soundproof barriers were erected around the patient to ensure the quality of the recordings, aiming to isolate them from noise as much as possible (Fig. 1a).

#### Preprocessing of audio signal

Initially, all audio recordings were manually audibly reviewed to identify and exclude samples with significant issues affecting recording quality, such as noticeable noise interference and character pronunciation errors. Correspondingly, the neural signals associated with these excluded audio samples were also removed. For training and evaluating the model, the audio data must be labeled, including the start and end points of each character’s pronunciation, as well as the segmentation points of the initial and final sounds during the pronunciation process of each character. In this study, we adopted a combination of automatic labeling using software and expert annotation. All audio materials were first automatically labeled using the PRAAT software^45^, after which all labeling results were verified and finally confirmed by three phonetics experts.

### Analysis of audio signal

#### Acoustic feature extraction

In the design process of the three element prediction models, relevant acoustic features were utilized, to enhance the prediction performance. Specifically, the initial prediction model employed acoustic-phonetic features, which are attributes inherent to the initials themselves, rather than being related to the recorded audio signals. Chinese is a tonal language, where speakers can modulate the pitch of their voice by adjusting the tension of their vocal cords to produce variations in pitch height and contour, thereby distinguishing different tones to convey various meanings. In this study, the pitch feature F0 was extracted from the audio signal and applied to the tone prediction model. The pronunciation characteristics of the final sounds can be assessed through three formants, with the first two formants being related to the position of the tongue, making them suitable for use in the final prediction model. Parselmouth^46^, a Python implementation of PRAAT was applied to obtain pitch and formant data^45^.

#### Audio signal classification through support vector machine

In Figure 3, various results of audio signal classification are presented, where panel b shows the outcomes of tone classification, panel d illustrates the classification of monophthongs, panel e displays the classification results for all vowels, and panel g reveals the classification outcomes for final clusters. All classifications were conducted using a one-against-all multi-class support vector machine (SVM) algorithm. The following decision function is employed for classifying 𝒙_*new*_:

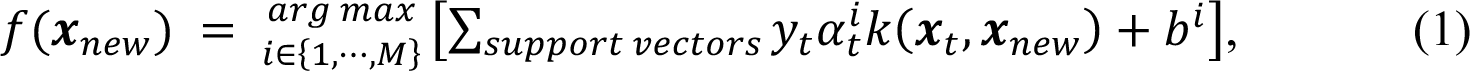

where, the 𝑡 represents the indexed sample, and 𝑖 ∈ {1, ⋯, 𝑀}denotes the total of 𝑀 binary Support Vector Machines (SVMs) that need to be trained, involving the solution of 𝑀 quadratic programming problems.

### Analysis of neural signals

#### Neural signals similarity analysis

Figure 2b displays the Pearson correlation between specific initials and the average of their corresponding category. We discuss two scenarios based on whether the initial is included in the corresponding category. In the first scenario, when the 𝑖^th^ initial does not belong to the 𝑗^th^ category of initials, we first calculate the average neural signal all trials of the 𝑖^th^ initial and the average neural signal all initials within the 𝑗^th^ category. The calculation of the correlation *R^i^_j_* is as follows:

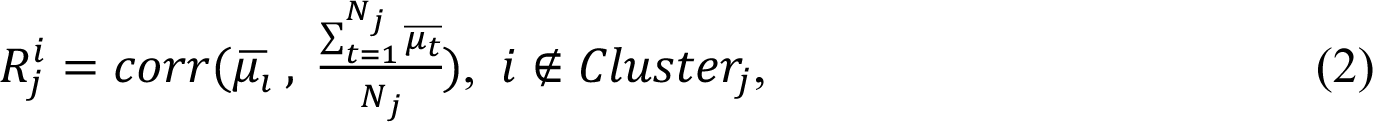

where, 𝑐𝑜𝑟𝑟 represents the Pearson correlation between two sequences, and 𝑁_;_ denotes the number of initials involved in the 𝑗^th^ category. In the second scenario, where the 𝑖^th^ initial belongs to the 𝑗^th^ category of initials, to exclude the positive bias introduced by self-correlation, we randomly divide the 𝑁 trials of the 𝑖^th^ initial into two groups and calculate their average neural signals, 𝜇̅ and 𝜇G, respectively. Then, we calculate the average of 𝜇̅ for all initials within the 𝑗^th^ category. The calculation of the correlation *R^i^_j_* is as follows:

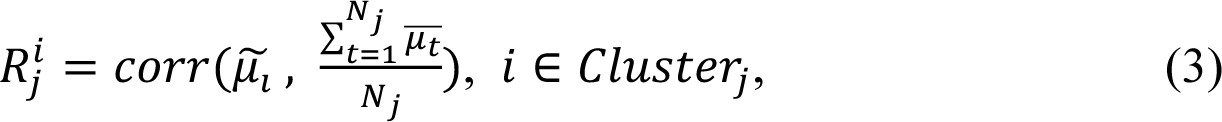

#### Clustering of finals

We cluster similar finals together due to the difficulty of differentiation brought about by dynamic nature of compound finals articulation. We consider a total of cluster number within the range of 10 to 15 to balance the granularity of classification and the classification accuracy. We employed the k-means clustering algorithm with silhouette coefficient to determine the optimal cluster number using the averaged formant features (F1, F2 and F3) across all audio samples of each final class as input, finding the optimal cluster number of 11. We then merge the finals within one cluster as one final cluster.

### Decoder performance evaluation

#### Character error rate (CER)

The character error rate (CER) is adopted as a metric to evaluate the performance of our proposed decoder, which is defined as follows:

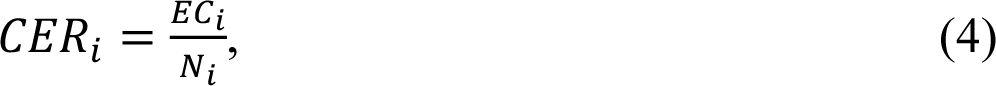

where *CER_i_* represents the character error rate for the *i^th^* evaluating sample, *EC_i_* denotes the number of characters in the *i*^th^ decoded result that do not match the ground truth, and 𝑁_*i*_ indicates the total number of characters in the *i*^th^ evaluating sample. Naturally, the character-level accuracy (CLA) is defined as:

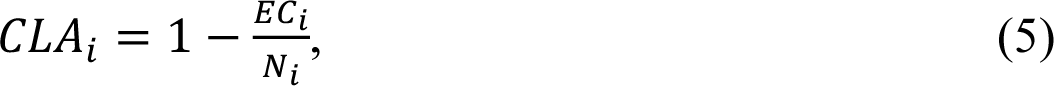

#### Syllable-level accuracy

During the evaluation of the performance of language model, syllable-level accuracy (SLA) is used to assess the model’s ability to convert pinyin syllables into Chinese characters. SLA is defined as:

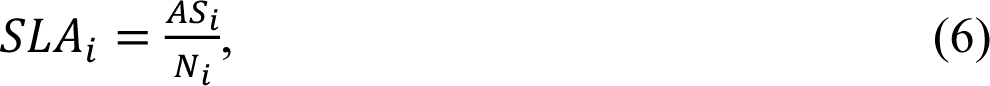

where *SLA_i_* represents the syllable-level accuracy for the *i*^th^ evaluating sample, *AS_i_* denotes the number of syllables in the *i*^th^ decoded result that match the ground truth, and *N_i_* indicates the total number of characters in the *i*^th^ evaluating sample.

#### Element-level accuracy

We adopt element-level accuracy (ELA) to evaluate the performance of the language model since it reflects the language model’s elements correction capability within a syllable. First, each Chinese character in the decoded word or sentence is converted into a *Pinyin* syllable. Then, the initial, tone, and final components of each decoded *Pinyin* syllable are extracted. We calculate ELA as follows:

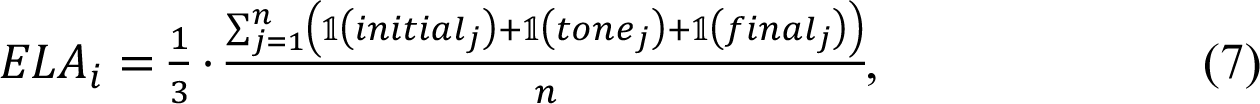

where *ELA_i_* represents the element-level accuracy for the *i*^th^ evaluating sample, 𝑛 represents the total number of characters of the *i*^th^ evaluating sample, 𝟙(⋅) is an indicator function with a value of 1 for a correct prediction of the corresponding syllable element and otherwise 0.

### Syllable elements prediction

#### Network architecture

We adopt a simple yet effective plain convolutional neural network (CNN) architecture as the backbone for syllable elements feature extraction. The detailed architecture of the network is provided in Supplementary Table 3. The subjects’ pre-processed sEEG of input channels 𝐶 is first passed to a stem layer containing a 1-D convolution layer. The 1-D convolution layer adopts a kernel size of 𝐾 × 1, a stride of 𝑆, and an output channel number of 64. The numbers of the kernel size 𝐾 and stride of 𝑆 are adjusted for each subject. Each convolution block consists of a 1-D convolution layer (followed by a batch normalization layer and a ReLU activation) and a max pooling layer. The number of the output channel of the first two blocks is set to 128, while the number of the last two blocks is set to 256. Then, we adopt a global average pooling layer and a fully connected layer to map the output of the last block to fit the prediction targets of different syllable elements.

#### Initial prediction model

We utilize the same CNN architecture (Network architecture part in Method) as the backbone for initial acoustic-phonetic feature prediction. On top of the learned hidden representation, a 2-layer multilayer perceptron (MLP) prediction head with a hidden dimension of 256 is utilized to predict the label of each acoustic-phonetic features of the initial. We follow the same experimental setup as described in Experimental setup part and report the mean and standard deviation of the top-1 accuracy for each acoustic-phonetic feature.

Inspired by the articulation of each Mandarin initial consonant, we proposed the articulation attribute integration (AAI) module to better capture the acoustic-phonetic features during the articulation process. The acoustic-phonetic features include the place of articulation (POA), manner of articulation (MOA), devoice, and aspiration. These acoustic-phonetic features can be considered as four attributes associated with each initial consonant. We first decompose the initial prediction problem into four attribute categorization subproblems. In particular, we utilize four learnable prototypes 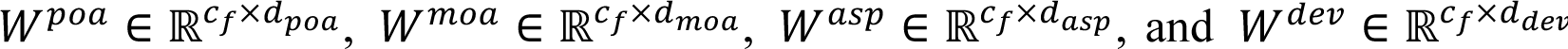, to map the neural features extracted by the CNN backbone to representations in the acoustic-phonetic attribute spaces. *c_f_* denotes the number of output channels of the CNN model, 𝑑_*poa*_, 𝑑_*moa*_, 𝑑_*asp*_, and 𝑑_*dev*_, represent the cardinality of the POA, MOA, devoice and aspiration attributes, respectively. Let 𝑧 ∈ ℝ^*c*^*f* be the extracted feature of the CNN; we concatenate the prototypes and perform attribute aggregation by multiplying with an adjacency matrix 𝔸̃:

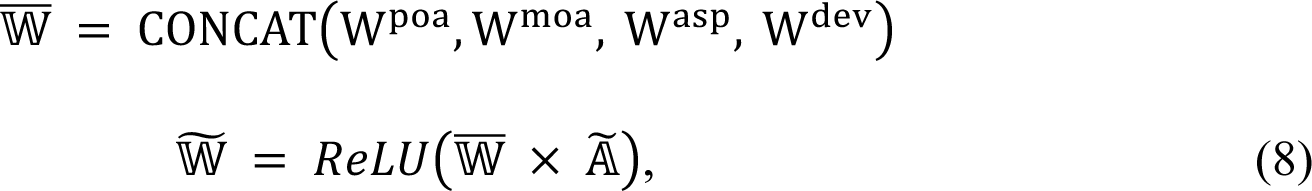

where 𝐶𝑂𝑁𝐶𝐴𝑇 denotes the concatenation operation and 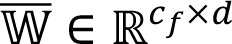 where *d* = 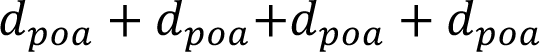. The adjacency matrix Ã is defined by adding self-loop as 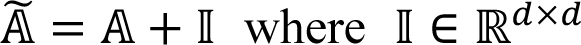 is the identity matrix. A_*i,j*_ = 1 if the attribute 𝑖 and attribute 𝑗 coexist in the onset process of any initial consonant and A_*i,j*_ = 0 otherwise. Notice that 𝔸 is only initialized by the previous condition and updated during training. The aggregated prototypes 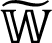 are then multiplied with feature vector 𝑧 to predict the four attribute labels. A 2-layer multilayer perceptron (MLP) prediction head with a hidden dimension of 256 is utilized to predict the initial label. The overall loss function is defined as:

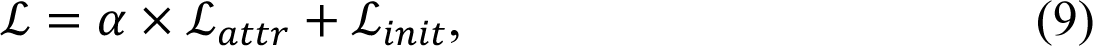

where *L_attr_* and *L_init_* stand for categorical cross-entropy loss for attribute classification and initial classification, respectively. 𝛼 is a scaler to balance the two losses.

#### Tone and final cluster prediction model

In the field of speech processing, the key factor in tone perception relies on pitch-related features where the fundamental frequency (F0) plays a crucial role. Likewise, each vowel is distinguished by a unique set of resonating frequencies of the vocal tract, referred to as formants. Formant features F1 and F2 correspond to the vertical and anteroposterior positions of the tongue, respectively, while Formant F3 indicates the tongue’s constriction in the palatal region. The similarity of formant features among different finals serves as a valuable indicator for measuring the articulation process, which we aim to decode using sEEG. Besides directly predicting the tone and final using the extracted from CNN, we introduce a neural-audio regularization (NAR) module, which enhances the CNN’s ability to model the similarity of the articulation process by using formant features as guidance. Let 𝑍 ∈ ℝ^*B×C*^ be the extracted feature from a training batch of sEEG data associated with the corresponding audio data, where 𝐵 denotes the batch size. It is worth noting that for tone prediction, we utilize the F0 feature, while for the final prediction, we use F1, F2, and F3. Subsequently, we calculate the pairwise Euclidean distance of 𝐷_*n*_ and 𝐷_*α*_ for both 𝑍 and 𝐹. Following this, we employ the Student’s t-distribution for similarity measurement:

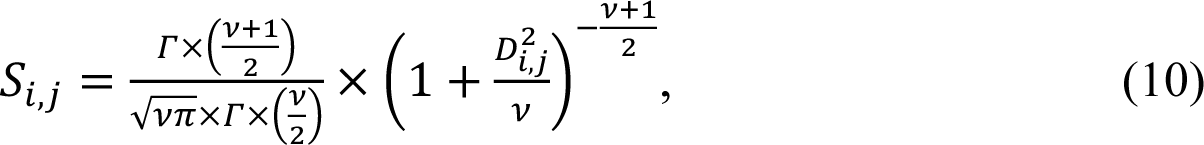

where 𝛤 and 𝜈 refer to the gamma function and the degree of freedom, respectively. 𝐷 denotes the Euclidean distance. The objective function to train NAR is defined to be the logistic loss between 𝑆_α_ and 𝑆_*n*_:

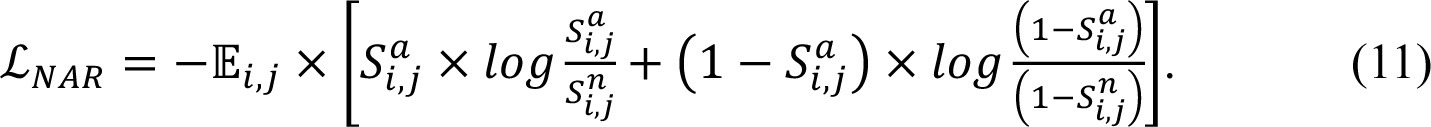

The overall loss function is defined to be a weighted sum of *L_NAR_* and the corresponding categorical cross-entropy loss for either tone or final prediction.

#### Experimental setup

The experiments were conducted using PyTorch^47^ on NVIDIA V100 GPUs. For investigating syllable elements level prediction, the data from three trials were combined and divided into 80% for training and 20% for evaluating, using 20 distinct random seeds. This setup facilitated the calculation of mean and variance for performance metrics. We carry out experiments on all four subjects with the same experimental setup. For each subject, 20% of the training data from the first repetition was allocated for tuning size 𝐾 and stride of 𝑆. The specific choices of kernel size and stride for each subject are detailed in Supplementary Table 4. The models were trained using the AdamW optimizer, with a base learning rate of 𝑙𝑟 = 0.0003 . Momentum parameters are set to 𝛽_-_ = 0.9, 𝛽_a_ = 0.009, and a cosine decay approach was adopted for the learning rate schedule. To prevent overfitting, a dropout rate of 0.3 was utilized. The training was conducted over 80 epochs, incorporating an early stopping criterion based on a five-epoch non-improvement threshold on the validation set. The complete list of parameters used for model training is available in Supplementary Table 5. For sentence-level performance assessment, syllable element prediction models were trained on the entire dataset for each subject over a fixed epoch number of 50.

#### Elements prediction with support vector machine

In this study, we adopted support vector machine (SVM) classifiers as baseline performance for syllable elements prediction. We follow the exact same data split strategy as described in Experimental setup part with the same evaluation metrics to ensure consistency across comparisons. For training of the SVM classifiers, we adopted the radial basis function (RBF) kernel with the regularization parameter and the kernel parameter gamma grid searched with five-fold cross validation on the training set for each random seed. We adopted the one-vs-the-rest (OvR) strategy for multiclass classification, as previously mentioned in **Audio signal classification through support vector machine part.**

#### Language model

Given a sequence of syllable elements predictions, denoted as 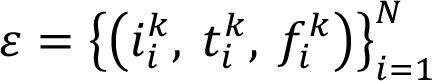 a language model is designed to output the most probable sentence where N represents the total number of characters in the sentence. The elements 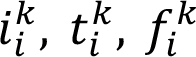 represent the top-k predictions for the initial, tone, and final elements of the *i^th^* character, respectively. The model’s top-3 accuracy for tone and initial predictions, as shown in Figure 3j and Figure 2f, reveals that the three most probable predictions often correctly identify these elements. Our approach involves constructing a 5-gram language model that evaluates the probability of initial and tone sequences, denoted as 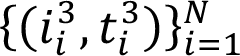, using the top three predictions for each. The training of this 5-gram model utilized CLUECorpus2020^48^, a comprehensive Mandarin Chinese corpus containing 35 billion Chinese characters retrieved from Common Crawl. CLUECorpus2020 contains a wide range of topics, including news, entertainment, sports, health, international affairs, movies, celebrities, and more. To develop our 5-gram model, we randomly selected 20% of this training set and implemented the model using KenLM^49^. Instead of employing a character-based vocabulary, we have constructed a specialized vocabulary comprising pairs of initials and tones. This approach results in a vocabulary of 84 distinct elements, derived from 21 initials combined with four different tones. We then utilize the Google Pinyin API to convert the initials from the top 50 probability sequences into Mandarin characters, forming sentences. For each initial sequence, we extract two sentence suggestions, thereby cumulatively creating a total of 100 sentences.

The majority of current Pinyin input methods do not rectify incorrect or illogical initial sequences. Instead, they endeavor to construct a sentence from the input, often resulting in combinations of characters that resemble sentences but lack coherent, natural language meaning. Consequently, it is advantageous to utilize large language models (LLMs) to determine the most probable sentence out of 100 potential options generated, harnessing the LLM’s capacity to discern coherent and meaningful language from a multitude of sentence possibilities. In our approach, we utilize the Chinese-LLaMA-7B model^50^ to evaluate the likelihood of different sentence options. Perplexity 𝒫𝒫ℒ of each sentence candidate is calculated using the following equation:

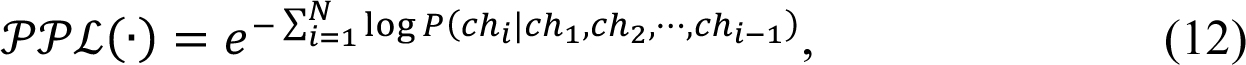

where 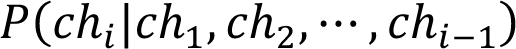 represents the probability assigned by the LLM to the character 𝑐ℎ_*_ based on the sequence of preceding characters. We then select the sentence with the lowest perplexity as the final output. This method ensures that the output is a grammatically correct sequence of characters and a coherent and contextually appropriate sentence.

### Contribution of different anatomical brain regions

We analyze the contribution of brain regions from two distinct perspectives. The first perspective utilizes sEEG signals from single functional brain regions as system inputs for Mandarin Chinese decoding, where the decoding performances of these regions serves as the metric for assessing their contribution level. The second perspective involves using all channels as model inputs, where we determine the contribution of brain regions based on the significance level of different channel signals in the decoding decision channel saliency calculated by class activation maps^51^.

#### The decoding accuracy of different brain regions

We train syllable element prediction models utilizing sEEG signals of channels that cover the specific brain regions that we aim to analyze and report the prediction performance. We follow the same experimental setup as described in Sec. 7.4 for syllable element prediction model and evaluations. This method allowed for a focused investigation of each region’s role in the decoding process from the perspective of the prediction performance. We follow the same experimental setup described in **Experimental setup part** for both evaluation approaches.

#### The contributions of electrodes located in different brain regions to decoding

Channel saliency score is calculated by class activation maps^51^. Given an sEEG signal 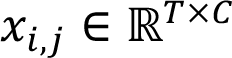 where 𝑇 is the length of the signal and 𝐶 is the total number of channels. We calculate a channel saliency score 𝜁:

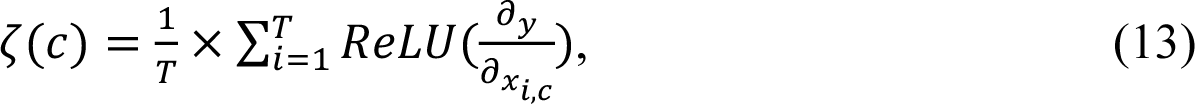

where 𝑦 is the one-hot logit vector corresponding to the true label of the underlying prediction task, and 𝑅𝑒𝐿𝑈 is the activation function. As illustrated in Equation 6, the partial derivative is calculated with respect to the input sEEG signal and averaged over the time dimension. This measure indicates how changes in the information content of channel c influence the accurate prediction of the true label. We then normalize the saliency score 𝜁(𝑐) for each channel 𝑐 to be between zero and one. The second measurement provided a comprehensive understanding of how the change of information from specific brain regions affects the Mandarin Chinese decoding process, highlighting the contribution level of each channel when using all-channel sEEG for language decoding.

### Model temporal saliency of syllable prediction models

Similar to the calculation of channel saliency score in Eq. (12), we calculate the temporal saliency score η(𝑡):

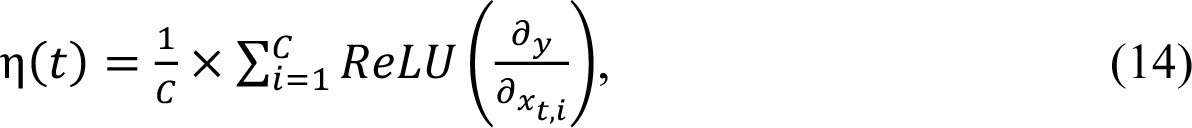

where 𝑥_2,*_ refers to sEEG signal of channel 𝑖 at time stamp 𝑡. Different from Eq. (13), the partial derivative is averaged over the channel dimension.

## Data availability

The data that support the findings of this study are available from the corresponding author upon reasonable request. Source data for figures are provided with the paper

## Code availability

All main code can be found on GitHub at https://github.com/diwu121/sEEG2Chinese.

## Extended Data

**Extended Data Table 1.**
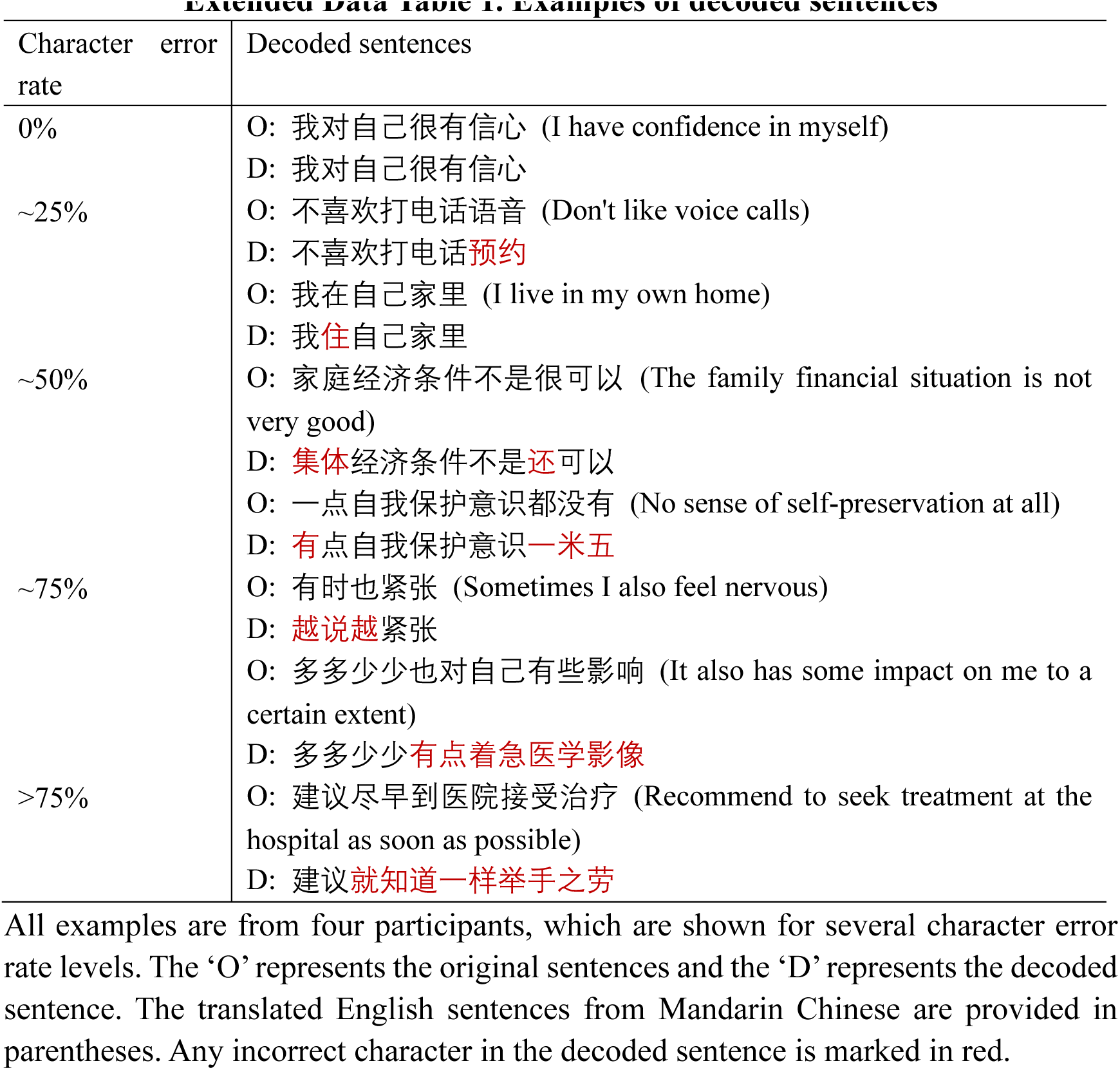
Examples of decoded sentences

**Extended Data Table 2.**
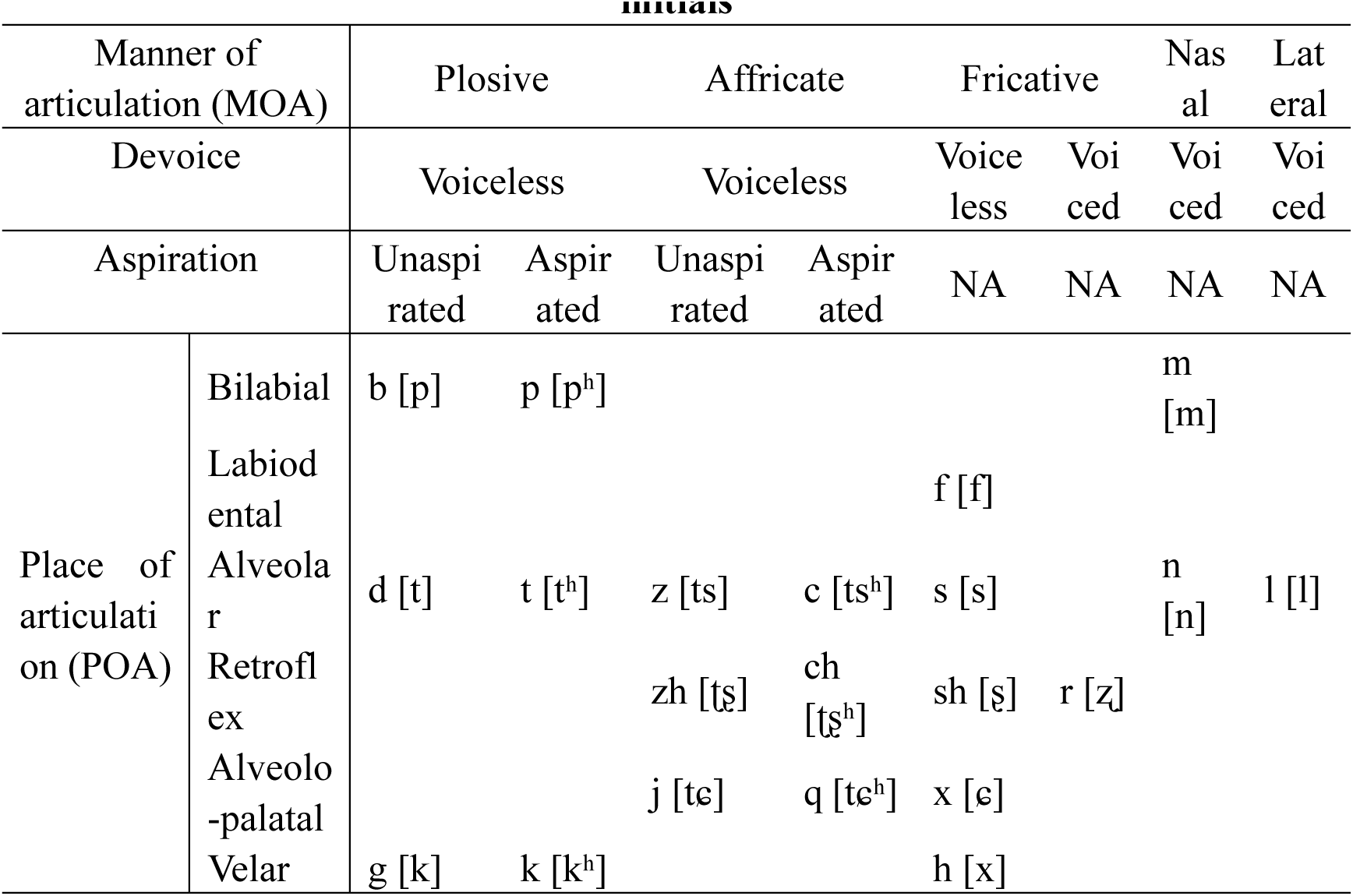
The articulation features of Mandarin Chinese *Pinyin* initials

**Extended Data Table 3:**
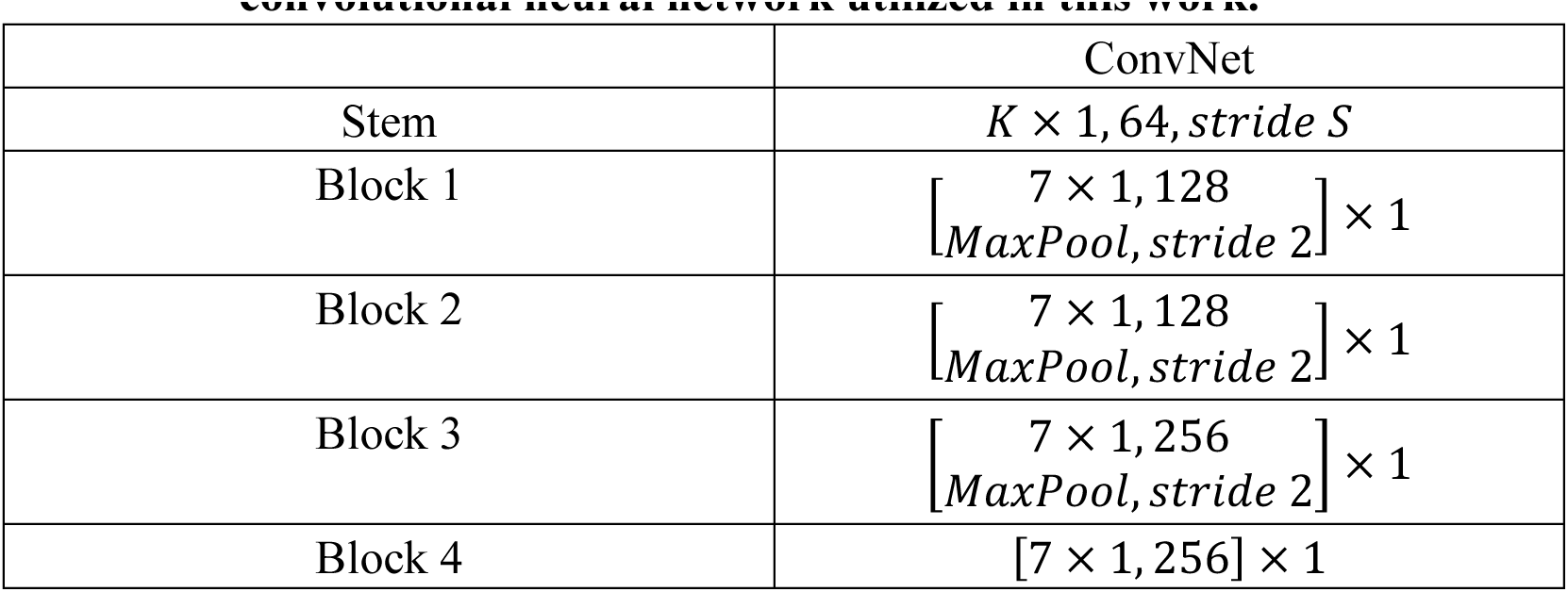
Detailed architecture specifications for the convolutional neural network utilized in this work.

**Extended Data Table 4:**
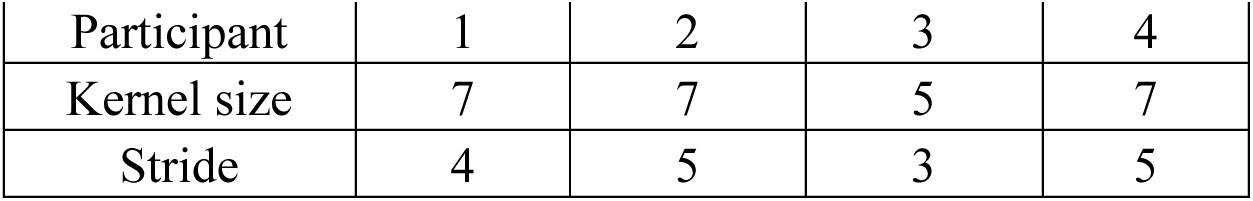
Architecture network and training parameters for each participant

**Extended Data Table 5:** Optimizing setup for syllable element prediction.

|  |  |
| --- | --- |
| Configuration | ConvNet |
| Training epochs | 80 |
| Optimizer | AdamW |
| Base learning rate | 0.0003 |
| Weight decay | 0.01 |
| Optimizer momentum | $\beta_1 = 0.9, \beta_2 = 0.009$ |
| Batch size | 32 |
| Learning rate schedule | cosine decay |
| Dropout ratio | 0.3 |

**Extended Data Fig. 1.**
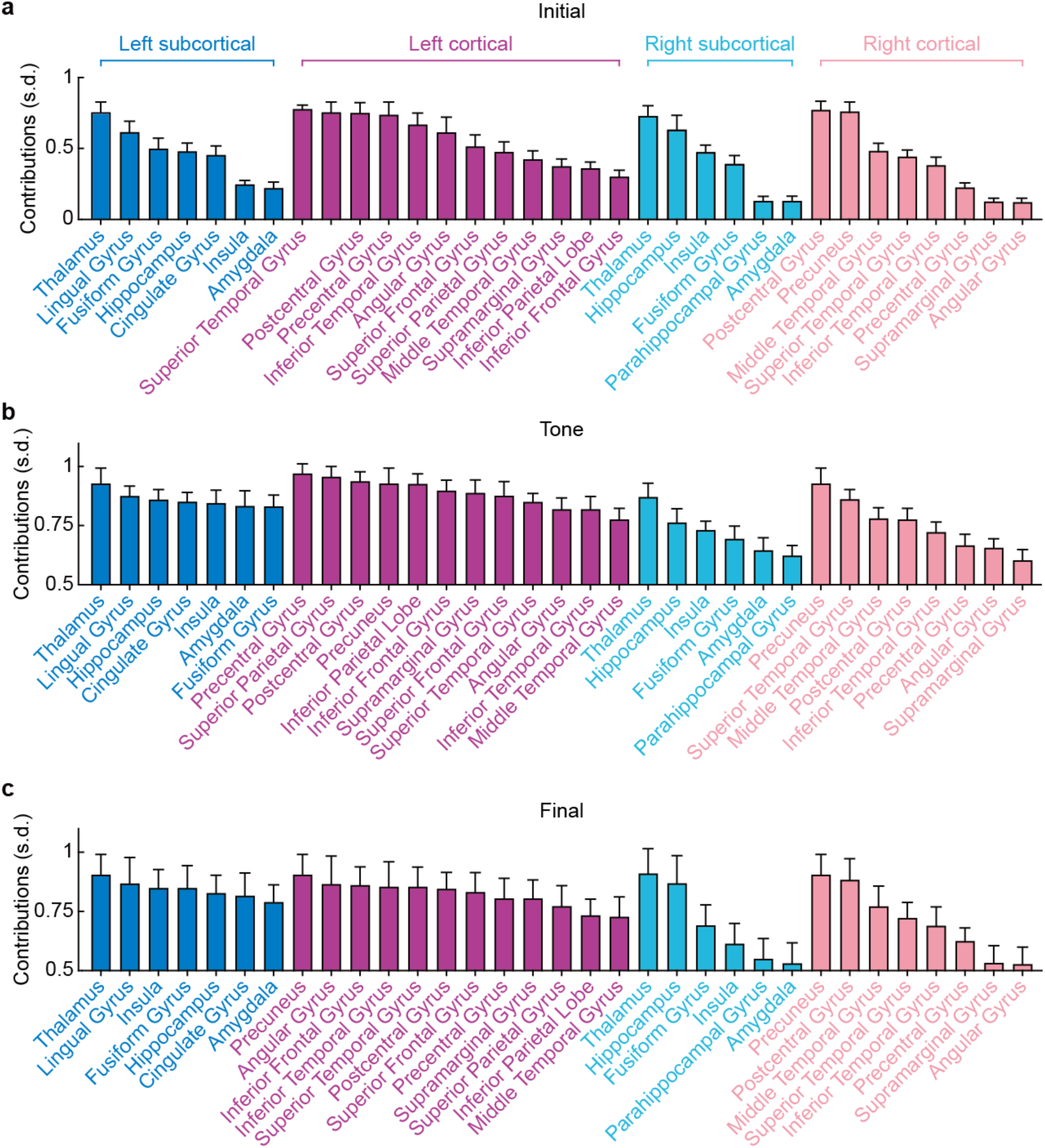
| The contributions of each anatomical area in cortical and subcortical structures of right and left hemispheres of the brain to predicting initials (a), tones (b), and finals (c), as measured by the gradient of the loss function with respect to the input data. The contributions of different participants to the same anatomical area are grouped together.

**Extended Data Fig. 2.**
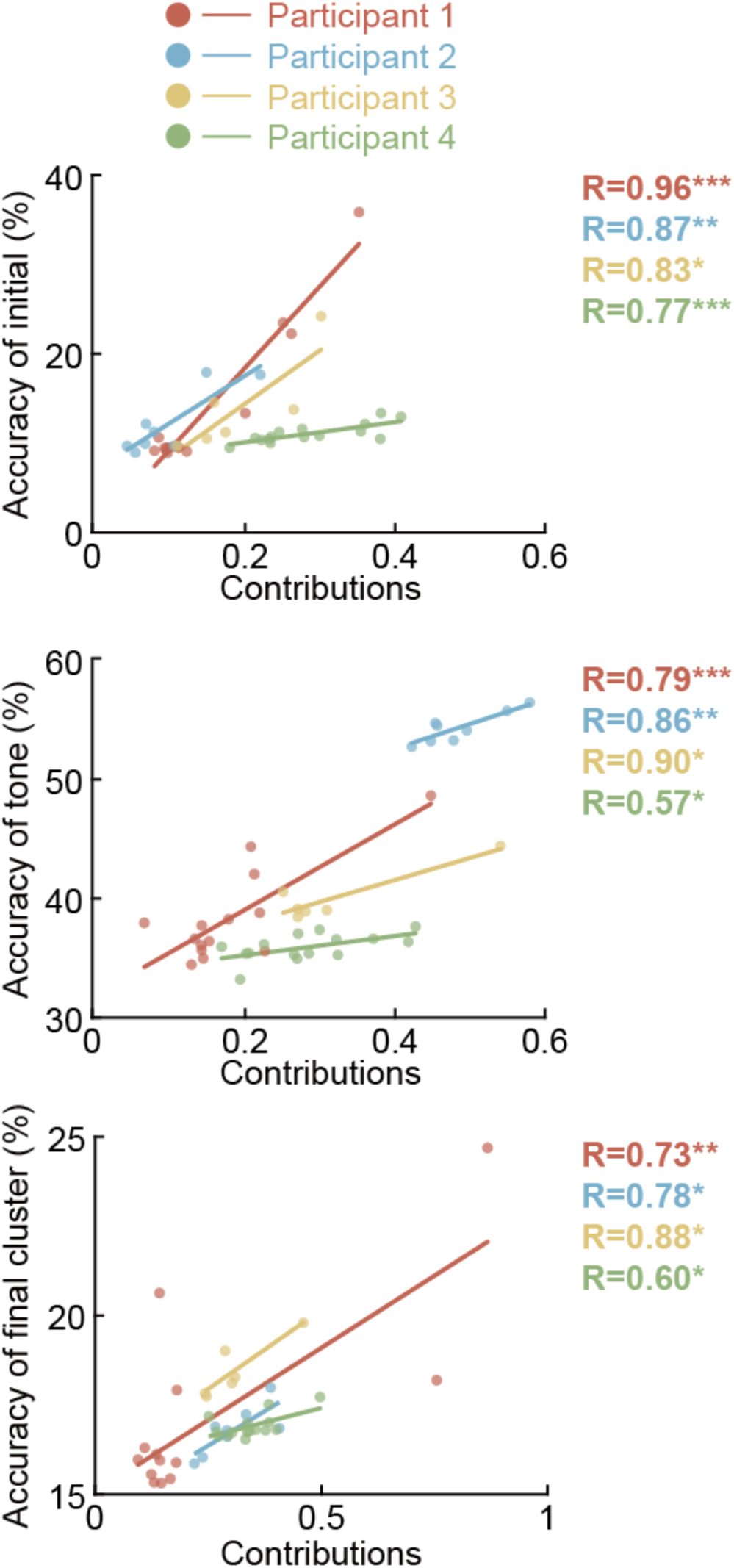
| The correlation between prediction accuracies and the corresponding contributions of each anatomical area in three syllable elements prediction models. The Pearson’s correlation coefficient (R) of three syllable elements is calculated separately, with different colors representing different participants (corresponding to Fig. 1). The contributions are measured by the gradient of the loss function with respect to the input data. Each dot represents a brain region’s contribution and decoding accuracy for prediction models of initial, tone, and final cluster. \**P* < 0.05; \*\**P* < 0.01; \*\*\**P* < 0.0001.

**Extended Data Fig. 3.**
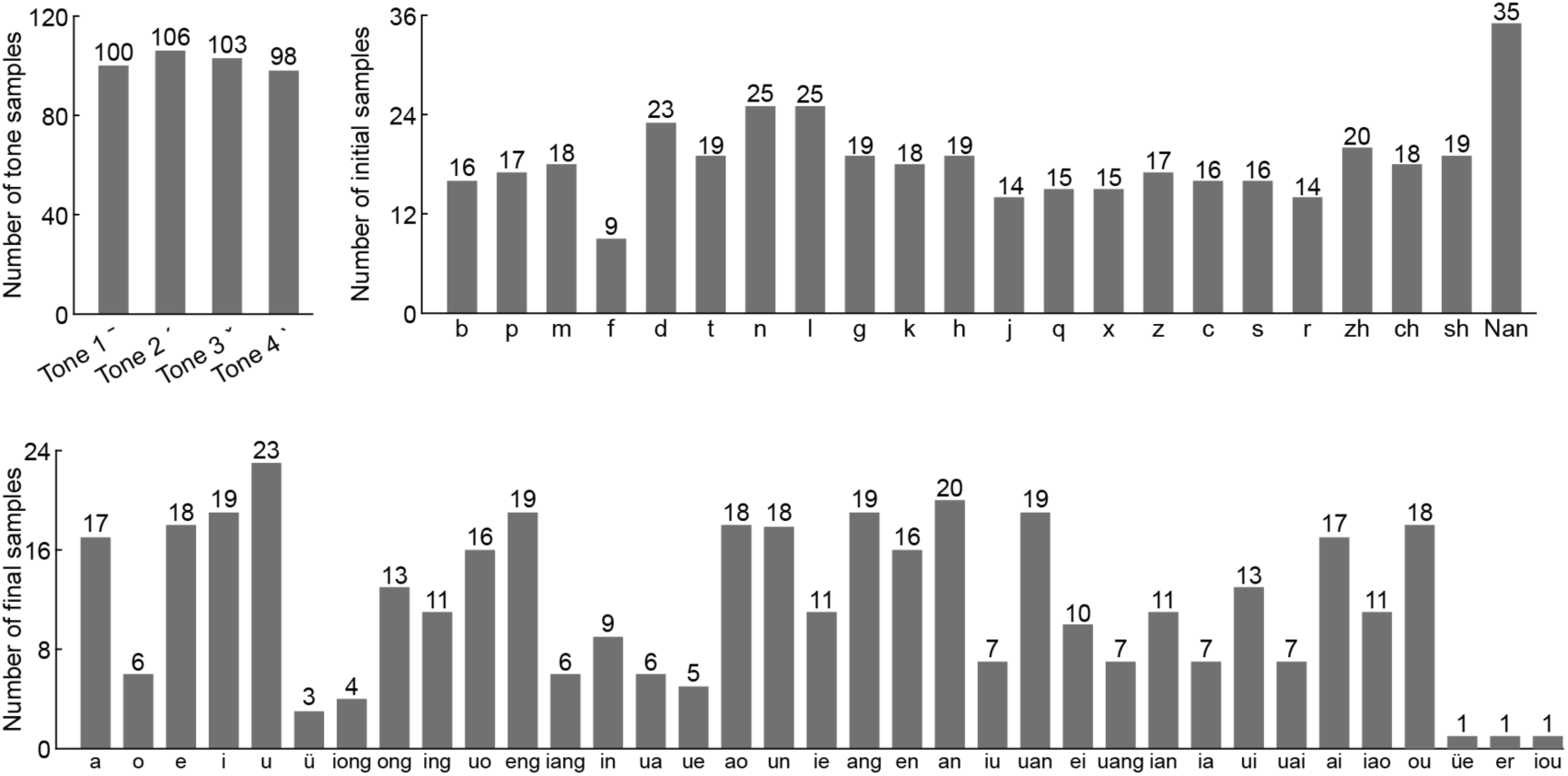
| Each syllable consists of a tone, a final (rhyme), and at most one initial (with some syllables lacking an initial). The charts display the distribution of the number of initials, tones, and vowels across the 407 syllables (the experimental tasks used for training the three element prediction models).

